# More than expected: the abundance of yellow-legged gulls *Larus michahellis* breeding in the historic centre of Venice and the initial effects of the new waste collection policy on the population

**DOI:** 10.1101/2021.01.01.425039

**Authors:** Francesca Coccon, Lorenzo Vanni, Caterina Dabalà, Dimitri Giunchi

## Abstract

Yellow-legged gull, *Larus michahellis*, has undergone widespread colonization of the urban environment in recent past. Starting in 2000 this affected the historical centre of Venice, 24 roof-nesting pairs being recorded in 2005, with this number increasing significantly in the last decade. In 2016, the waste management company of Venice established a new door to door garbage collection system to prevent the accumulation of rubbish in the streets and limit the trophic resources available for the species. The study provides an up-to-date estimate of the urban population of yellow-legged gulls, using Distance Sampling method. We also studied the effect of the new system on the species by comparing the population estimate before and after the change and by analysing the trend of individuals collected in the old town by the service of wildlife recovery during 2010-2018. Results estimate 440 breeding pairs (95s% confidence interval: 326-593) in June 2018 and show a 34% decrease of breeding pairs in 2018 with respect to 2017, as well as a decrease in number of 1-year birds and *pulli* collected by wildlife recovery service starting from 2016, year of the policy implementation. Our data did not show a significant decrease in the overall number of individuals, suggesting the new policy has a stronger effect on the breeding success of the species than on adult survival. This study emphasizes the importance of preventing rubbish accumulation in the streets as factor for reducing the abundance of urban yellow-legged gulls.

## Introduction

Over the centuries and particularly from the beginning of the industrial revolution, man has profoundly transformed the Earth’s surface by converting the original landscape into anthropic ecosystems, mainly represented by urban areas and farm fields (Berry 2008; Houghton 1994; Meyer and Turner 1992), thus generating major pressures on other life forms, exerting selective pressures driving the evolution of many species (Alberti et al. 2003; Albuquerque et al. 2018; Ellis 2015). As a whole, human actions and urban sprawl have led to a trivialization and homogenization of landscapes by destroying and fragmenting the original habitats (McKinney 2006). This has resulted in a generalization of the wildlife communities (Marzluff 2001), with the loss of rare and specialized species (commonly known as ‘losers’) and an increase of generalist and adaptable ones (‘winners’) (McKinney 2008; McKinney and Lockwood 1999). The latter species, defined as synanthropic (from the Greek syn + anthropos, together with man), have adapted to live in highly anthropized habitats, well tolerating the disturbance effects from anthropogenic pressure and activities (e.g. traffic noise, air, water and soil pollution) and indeed taking advantage of human presence (Rodewald and Shustack 2008).

Among birds, gulls (*Larus* spp.) become so well adapted to the urban context that they have become superabundant and have started to be considered a pest species (Blokpoel and Spaans 1991; Feare 1991). In particular, the yellow-legged gull, *Larus michahellis*, has undergone a widespread population explosion over the past 30 years in the Mediterranean basin (Vidal et al. 1998). In Italy, the species showed a clear demographic increase during the second half of the 1900s, passing from 24.000-27.000 breeding pairs in 1983 (Meschini and Frugis 1993) to the 45.000-60.000 in the early 2000s (Brichetti and Fracasso 2006). The demographic increase of the species has been accompanied by a spread of the breeding range with the colonization of the urban environment, where gulls began to nest on the rooftops and terraces of buildings (Monaghan and Coulson 1977). The urbanization of gulls has led to a series of problems: from the acoustic nuisance, especially in the reproductive period, to fouling and damaging of the architectural and monumental heritage, to the aggression of adults in defence of their chicks (Dwyer et al. 1996; Soldatini et al. 2008) and the conflicts with commercial premises such as fish markets, butchers, bars or street food vendors (Belant 1997; Serra et al. 2016). In Italy, the first urban colony of yellow-legged gull dates back to 1971 in Rome, but it is only since the early 1980s that the phenomenon of gulls nesting on roofs has increased and spread to other cities: from Sanremo (1982), to Livorno (1984), Genova (1986), Trieste (1987), Naples (1990), highlighting a rapidly growing trend (Fraissinet 2015).

In the historical city centre of Venice the first individuals showing reproductive behaviour were observed in 2000, in 2005 24 roof-nesting pairs were recorded in the whole urban area (Soldatini and Mainardi 2006), while the latest published estimate available for the species indicate 50 pairs breeding in the old town (Bon and Stival 2013). However, over the last 25 years, the breeding population of yellow-legged gulls has increased dramatically in the lagoon surrounding the city of Venice, rising from an estimated average number of 1350 pairs in 1990-92 (Scarton 2017) to the 4803 in 2013-2015 (Scarton and Valle 2017). The demographic growth highlighted by the species in the Venetian lagoon was followed by an increase in the number of individuals attending the historic centre of Venice both for breeding and feeding purposes, exacerbating the problems of coexistence with this species (Coccon et al. 2019). Among the main critical issues which have been widely documented by the local press, are the recurring attacks on passer-by to steal their food and the breaking open of garbage bags dropped by residents and tourists on the street and spreading of their contents both on the ground and in the city’s canals (Coccon and Fano 2020). The latter situation has created what appear to be open-air landfills in the city leading to a negative image of Venice, considered an iconic place all over the world. To counter such problems, the public waste management company of Venice (Veritas Spa) established a new door to door garbage collection system and garbage self-disposal to temporary waste disposal stations located on boats moored in specific areas of the city. This was to prevent the accumulation of rubbish in the streets and limit the amount of trophic resources available for the yellow-legged gulls. With this regard, a recent study (Coccon and Fano 2020) revealed that the new urban waste collection regime had a significant effect on lowering both the presence of waste and gulls attending the city for feeding purposes. However, the effect of the new policy on yellow-legged gulls’ breeding population has not been investigated yet. To fill this gap, we started a monitoring program with the aim of providing an updated and accurate estimate of the species (i.e. number of individuals and breeding pairs) in the historic centre of Venice. This is urgently needed since data from the last monitoring of the species in the city dates back to 2005 and are no longer usable to support decision-making concerning urban control and management of the species. To achieve this goal, we used a Distance Sampling approach (Buckland et al. 2001) applied to counts performed from elevated observation points. As far as we know (see Ross et al. 2016 for a review), Distance Sampling has never been used before for monitoring the target species in an urban environment, although it is note that such method produces estimates of population more precise than direct counts (Coulson and Coulson 2015). We also investigated the initial effects of the new waste collection system on the yellow-legged gulls using two different approaches: *i*) by comparing the abundance of individuals and breeding pairs recorded in a given area of the city before and after the waste policy change; and *ii*) by analysing the trend of individuals (i.e. *pulli*, juveniles and adults injured or in difficulty) collected in the urban area from 2010 to 2018 by the service of wildlife recovery performed by the Metropolitan City of Venice, as we assumed this trend may be used as an index for the urban gulls’ population for those years in which monitoring was not conducted.

Results from this study will contribute to updated and accurate knowledge of the situation of yellow-legged gulls in Venice and will provide useful information on whether or not the new waste collection policy has been successful in lowering the problem of urban gulls. Finally, our work emphasises the potential value of Distance Sampling to provide accurate estimates of population and the possibility for this method of being applied in other urban contexts for monitoring and management programs of the species in the medium and long run.

## Methods

### Study area

Our study has been conducted in the historic centre of Venice, Italy (45°26′13.67″ N 12°19′57.54″ E, 6.52 km^2^), which is divided into six districts (Figure 1): Dorsoduro (0.97 km^2^), Santa Croce (1.42 km^2^), San Polo (0.34 km^2^), San Marco (0.54 km^2^), Cannaregio (1.40 km^2^) and Castello (1.90 km^2^). Among them, Cannaregio is the most populous with 15605 inhabitants, followed by Castello (13424 inhabitants), Dorsoduro (6430 inhabitants), Santa Croce (4939), San Polo (4612) and San Marco (3750) (data updated 1^st^ January 2017 provided by the Statistics and Research Office of the Municipality of Venice). With regards to the latter, despite the particularly low residential rate, there is the highest production of waste (amount of waste produced daily in the district) which is equal to 32 tons per day (data updated 31^th^ December 2017 provided by the public waste management company of Venice, Veritas). Tourist pressure contributes significantly to this value, as demonstrated by the conspicuous number of B&B and tourist accommodation facilities officially declared to the Municipality of Venice, present here (n= 199) (see Table S1).

**Fig. 1.**
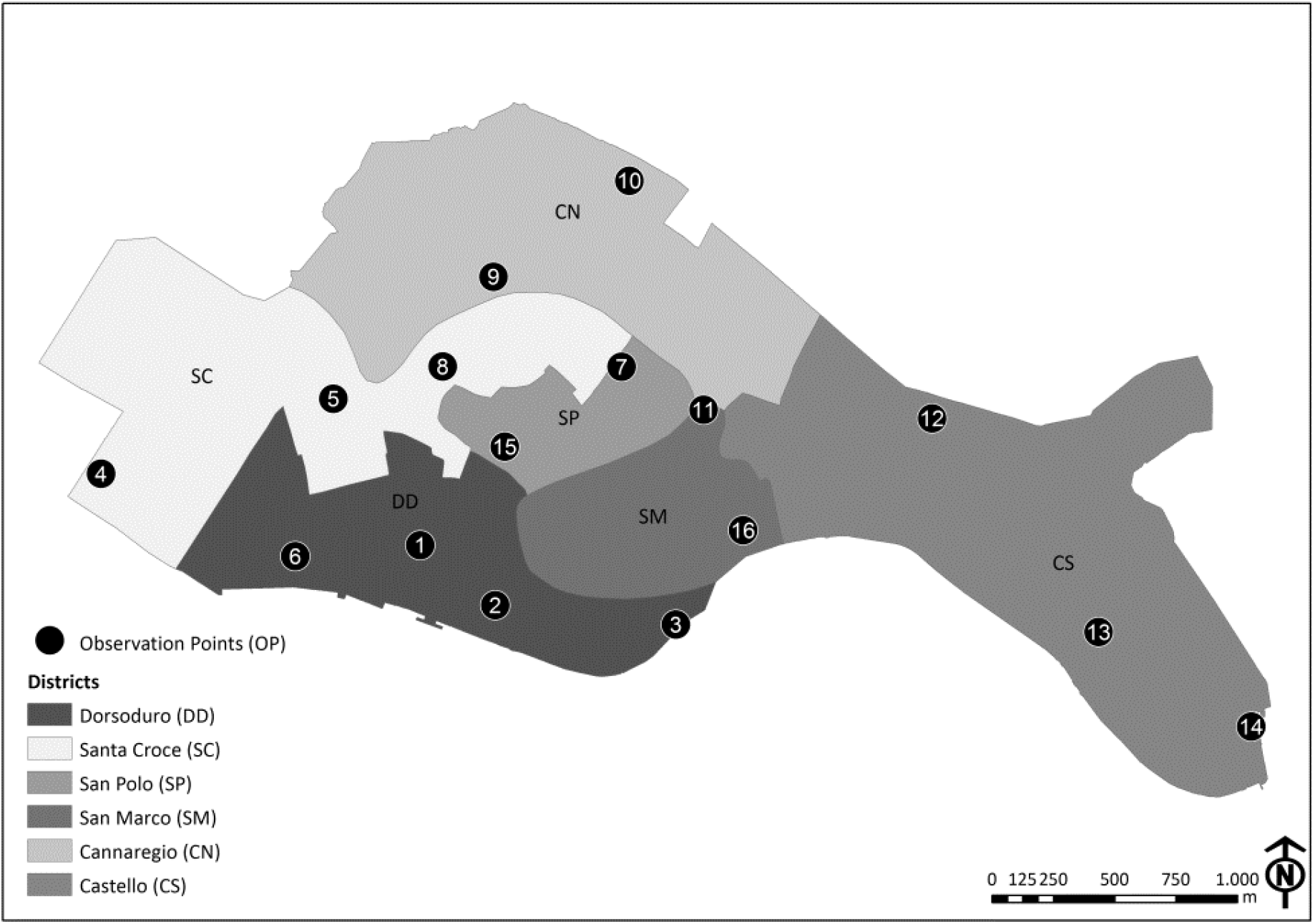
Historic centre of Venice divided into six districts: Dorsoduro (DD), Santa Croce (SC), San Polo (SP), San Marco (SM), Cannaregio (CN) and Castello (CS). The observation points (numbered black dots) selected in each district for monitoring yellow-legged gulls are also depicted (see Table S2 for specific information on observation points)

### Data collection

The monitoring activity was performed from 16 observation points (OPs, bell towers or private and public panoramic terraces; Table S2) distributed in all the six above-mentioned districts (Figure 1). The number and distribution of OPs in each district depended on the availability of vantage points and on the extent and visibility (height of the nearby buildings relative to the OP height) recorded in the area. A pilot survey was carried out in 2017 focusing on Dorsoduro and Santa Croce districts (OP 1 to 8 in Figure 1). In 2018 monitoring was extended to the whole historic centre of Venice, thus using all 16 OPs. In each survey, we performed two monitoring sessions: in March, at the beginning of the reproductive season, when yellow-legged gulls start to occupy their reproductive sites and in June, when the breeding season reaches its climax and the probability of detecting juveniles near the nest is higher (Fracasso et al. 2011). The monitoring activity was performed in the morning, starting at dawn to detect the birds’ peak activity time and ending at most 3-4 hours after dawn (Parra-Torres et al. 2020). This period was necessary in order to explore the roofs in detail and to detect the individuals. During surveys we recorded all the yellow-legged gulls detected while resting on the building roofs or on the ground, as well as those individuals that passed from flight to rest and vice versa. Data were collected by two professional ornithologists equipped with 10×42 binoculars (models: Leika Ultravid and Zeiss Terra ED 42) and one 20x-60x spotting scope (Kowa TSN-883 Fluorite). In this way, while one observer was intent on determining the position of the yellow-legged gulls, the other controlled their movements in order to avoid the possibility of detection errors such as missing of individuals or double counts. The number of breeding pairs was also recorded. We considered a breeding pair to be two adults observed in a suitable nesting habitat, showing a territorial behaviour and/or performing alarm calls, or one individual while incubating eggs or bringing food to the chicks. Even the sole presence of chicks at the nest was considered to be an indicator of a breeding pair. For data collection we used an application for mobile devices, specifically developed for this study, which allowed us to record the spatial location and the characteristics of each observation (no. of individuals, presence of a breeding pair, a nest or chicks). This was possible by using Google Satellite as the basic map for recording the data, with the possibility of switching to the high-resolution georeferenced orthophotos of the historic centre of Venice, provided by the Venice Municipality, if the internet connection was not available. Data were stored in the device in real time and then exported to a Geographical Information System (ESRI, ArcGIS 10.2 for Desktop) platform for subsequent processing. Individuals detected in clusters were treated as single sightings, geolocating the centroid of the group.

### Estimate of the abundance yellow-legged gulls’ breeding population

Data were analysed using distance sampling (Buckland et al. 2001; Buckland et al. 2015) which has been suggested as an effective bird survey method in urban environment (Giunchi et al. 2007).

In order to obtain reliable estimates using distance sampling, four assumptions need to be and were satisfied:

- Observation points need to be randomly distributed with respect to the species’ distribution. In our case the OPs were distributed opportunistically and not randomly, thus possibly leading to a biased estimate of population density. Given the peculiarity of the OPs, this bias could not be avoided, but it seems reasonable to assume that its effect on the abundance estimation should not be significant as the OPs was chosen independently from yellow-legged gull distribution.
- All yellow-legged gulls located on or very near the OP should be detected. Due to the characteristics of the OP (see Table S2), it was sometimes impossible to see directly below them and thus a blind area existed exactly below the OP. For this reasons all data were left-truncated at 10 m from the OP.
- Yellow-legged gulls were detected in their initial position, before being disturbed by the observer. To reduce this effect, monitoring started about ten minutes after the observers had taken position to allow yellow-legged gulls to get used to them.
- Distances were measured accurately. Thanks to the use of the mobile application for mapping all detected birds, distance from the OPs to each record were derived, using ArcGIS, with an estimated accuracy of ± 1 m. Distances were calculated by disregarding the height of the OPs and of the detected birds.

Distance data were analysed using the package Distance 1.0.1 (Miller et al. 2019) in the R 4.0.2 environment (R Core Team 2020). We performed two different types of analysis: in the first one we estimated the abundance of individuals, while in the second one we estimated that of breeding pairs, thus excluding all birds with no indication of reproductive activity (see description above). The data from each survey were analysed independently as the sample size was large enough (> 75 detections, Buckland et al. 2001) and preliminary data explorations indicated that the use of pooled data would not represent a significant improvement for the analysis (analysis not reported).

For the individual-based analysis we modelled the detection-probability function considering the clusters of individuals; density estimation was then calculated by multiplying clusters density by mean cluster size, since a preliminary check of the data did not indicate any size bias problem (Buckland et al. 2001). In the second analysis, the detection-probability function was modelled by considering one breeding pair as a single detection. In modelling the detection-probability function we considered four models (Thomas et al. 2010): half-normal key with cosine and Hermite polynomial adjustments, uniform key with cosine adjustments, and hazard-rate key with simple polynomial adjustments. To improve the fit, data were right-truncated at 500 m. As mentioned above, all data were also left-truncated at 10 m from the OP in order to take into consideration the blind area exactly below each OP. Mean cluster size was calculated using truncated data. For each OP the survey effort was obtained by considering the proportion of urbanized area included in the buffer calculated at the right truncation distance. In this way we excluded the area occupied by the water of the lagoon, which was not considered in the survey. The best model for the detection function in each survey was chosen using the Akaike Information Criterion, AIC (Buckland et al. 2001; Burnham and Anderson 2002). The selected model was then used to calculate density only if the Cramér-von-Mises goodness of fit tests were not significant (*P* > 0.05). Comparisons among parameters involved in these estimates were performed by considering the 95% confidence interval (95% CI), as suggested by Johnson (1999).

### Analysis of the initial effects of the new waste collection system on gull population

The new waste collection system was introduced in the districts of the historical city centre of Venice at different times. Specifically, starting from September 2015, it was applied on an experimental basis in a portion of Dorsoduro characterized by a low residential rate and then it was extended to the remaining part of this district in October 2016. The policy was then applied to Santa Croce and San Polo in March 2017, to San Marco in May and to Cannaregio in December. The last district was Castello, where the new method for collecting waste was introduced in May 2018 (Coccon and Fano, 2020). Due to funding limitation, it was not possible to estimate the abundance of the population of yellow-legged gulls before and after the implementation of the new policy in the whole city centre. For this reason, we decided to evaluate the initial effects of the policy change using two different proxies: *i*) we compared the density estimation of yellow-legged gulls in the two districts surveyed in both years 2017 and 2018 (i.e. Dorsoduro and Santa Croce), where the new policy has been implemented in 2016-2017; *ii*) we analyzed the trend of yellow-legged gulls collected in the Venice historical city centre from 2010 to 2018 by the service of wildlife recovery, as we assumed it may reflect the trend of yellow-legged gulls’ urban population for those year in which monitoring had not been conducted. In fact in a city like Venice, where you move by walking in the streets, it is particularly easy to notice if there are animals in difficulty, whether they are chicks, juveniles or adults, and it is possible to assume that most of them have been collected by the wildlife recovery service, both because of widespread interest of citizens in urban fauna, and efficiency of the service. Wildlife recovery data were provided by the Veneto Region-Direzione Agroambiente, Programmazione e Gestione ittica e faunistico venatoria-Unità Organizzativa Coordinamento gestione ittica e faunistico-venatoria Ambito Litoraneo-Sede Territoriale di Venezia.

#### Comparison of the density estimation before (2017) and after (2018) the change of the waste collection policy

Density estimates of individuals and breeding pairs in March and June 2017 and 2018 were obtained using the same approach as detailed above for population estimate, except for the use of the data belonging to only eight of the sixteen observation points (Ops from 1 to 8, Figure 1), which covered the two districts surveyed in both years (i.e. Dorsoduro and Santa Croce). We considered density and not abundance since the estimates concerned a fraction of the study area. We expected that the estimates in 2018 were noticeably lower than those obtained in 2017, especially concerning the number of breeding pairs, as an effect of the widespread reduction of food availability in 2018 following the implementation of the new waste collection system, as highlighted by Coccon and Fano (2020).

#### Analyses of the trend of yellow-legged gulls collected in the Venice historical city centre by the service of wildlife recovery from 2010 to 2018

The trend of yellow-legged gulls collected in the urban area by the service of wildlife recovery from 2010 to 2018 was analysed by means of a Generalized Additive Model (GAM) with Poisson error structure and log link estimated with the package *mgcv* 1.8-33 (Wood 2017) in the R 4.0.2 environment (R Core Team 2020). The predictors of the model were the year and the age class of the collected bird (two-level factor: a) > 1-year birds; b) 1-year birds or *pulli*). The model included the main effects “Year” and the ‘smooth-factor’ interaction “Year by Age”, which produced a separate smooth for the two levels of Age (Wood 2017). We used a thin plate regression spline as smooth basis, setting k = 8 as the maximum basis dimension; the actual degree of smoothing was estimated by general cross-validation (Wood 2017). Model assumptions (autocorrelation and distribution of residuals, homogeneity of variance, influential observation, overdispersion) were checked according to Wood (2017) using the *mgcv* package.

## Results

### Estimate of the abundance of yellow-legged gulls’ breeding population

Excluding the survey performed in June, only considering the breeding pairs, the half-normal key with no adjustment was selected for the detection function in all the analyses (Table 1, Figures S1 and S2). The fit of the models was good both for individuals and breeding pairs after a reasonable right-truncation which excluded ≤ 21.1% of observations from the analysis. According to the estimated detection probability, all models provided relatively comparable results in both the studied period.

**Table 1.** Ranking of candidate models considered in each survey performed in 2018 for estimating the abundance of individuals and breeding pairs of yellow-legged gull, based on the Akaike’s information criterion (AIC). K = Number of parameters; GOF (p) = p-value of the Cramer-von Mises goodness of fit test; 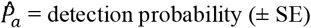; ΔAIC = difference in AIC from the best model.

| Survey | Model | K | GOF (p) | $\hat{p}_a \pm \text{SE}$ | ΔAIC |
| --- | --- | --- | --- | --- | --- |
| *Individuals* |  |  |  |  |  |
| March | Half-normal + Hermite polynomials | 1 | 0.97 | $0.42 \pm 0.02$ | 0.00 |
| Excluded observations: 20.7% | Half-normal + cosine | 1 | 0.97 | $0.42 \pm 0.02$ | 0.00 |
| | Uniform + cosine | 1 | 0.83 | $0.41 \pm 0.01$ | 2.61 |
| | Hazard-rate + simple polynomials | 3 | 1.00 | $0.41 \pm 0.06$ | 3.25 |
| June | Half-normal + Hermite polynomials | 1 | 0.70 | $0.46 \pm 0.02$ | 0.00 |
| Excluded observations: 21.1% | Half-normal + cosine | 1 | 0.70 | $0.46 \pm 0.02$ | 0.00 |
| | Hazard-rate + simple polynomials | 3 | 0.82 | $0.42 \pm 0.07$ | 1.82 |
| | Uniform + cosine | 1 | 0.54 | $0.44 \pm 0.02$ | 3.68 |
| *Breeding pairs* |  |  |  |  |  |
| March | Half-normal + Hermite polynomials | 1 | 0.77 | $0.39 \pm 0.03$ | 0.00 |
| Excluded observations: 15.6% | Half-normal + cosine | 1 | 0.77 | $0.39 \pm 0.03$ | 0.00 |
| | Uniform + cosine | 1 | 0.69 | $0.39 \pm 0.02$ | 1.52 |
| | Hazard-rate + simple polynomials | 3 | 0.97 | $0.33 \pm 0.08$ | 1.97 |
| June | Uniform + cosine | 1 | 0.73 | $0.42 \pm 0.03$ | 0.00 |
| Excluded observations: 17.3% | Half-normal + Hermite polynomials | 1 | 0.78 | $0.43 \pm 0.03$ | 0.56 |
| | Half-normal + cosine | 1 | 0.78 | $0.43 \pm 0.03$ | 0.56 |
| | Hazard-rate + simple polynomials | 2 | 0.63 | $0.46 \pm 0.06$ | 4.01 |

Density estimates, obtained using the best models selected according to AIC, turned out to be quite precise in both months of survey, with a coefficient of variation less than 0.15 in all cases (Table 2). The density of gulls was higher in March and then markedly decreased in June. According to our results the number of gulls in Venice varied on average between 2000 and 3000 individuals (Figure 2). It is important to emphasize that this estimate probably represents a minimum abundance as we did not consider birds in flight in our estimation.

**Table 2.** Density of individuals and breeding pairs of yellow-legged gull obtained for the whole city centre in each survey performed in 2018. Estimates were calculated using the best models chosen by the Akaike’s Information Criterion (AIC).

| **Survey** | **No. of observations**<br>**(clusters or pairs)** | **Density ± SE**<br>**(individuals or pairs/km<sup>2</sup>)** | **CV** |
| --- | --- | --- | --- |
| ***Individuals*** |  |  |  |
| March | 760 | 417.8 ± 45.9 | 0.11 |
| June | 779 | 312.2 ± 34.9 | 0.11 |
| ***Breeding pairs*** |  |  |  |
| March | 368 | 98.4 ± 14.3 | 0.14 |
| June | 272 | 67.4 ± 9.8 | 0.14 |

**Fig. 2.**
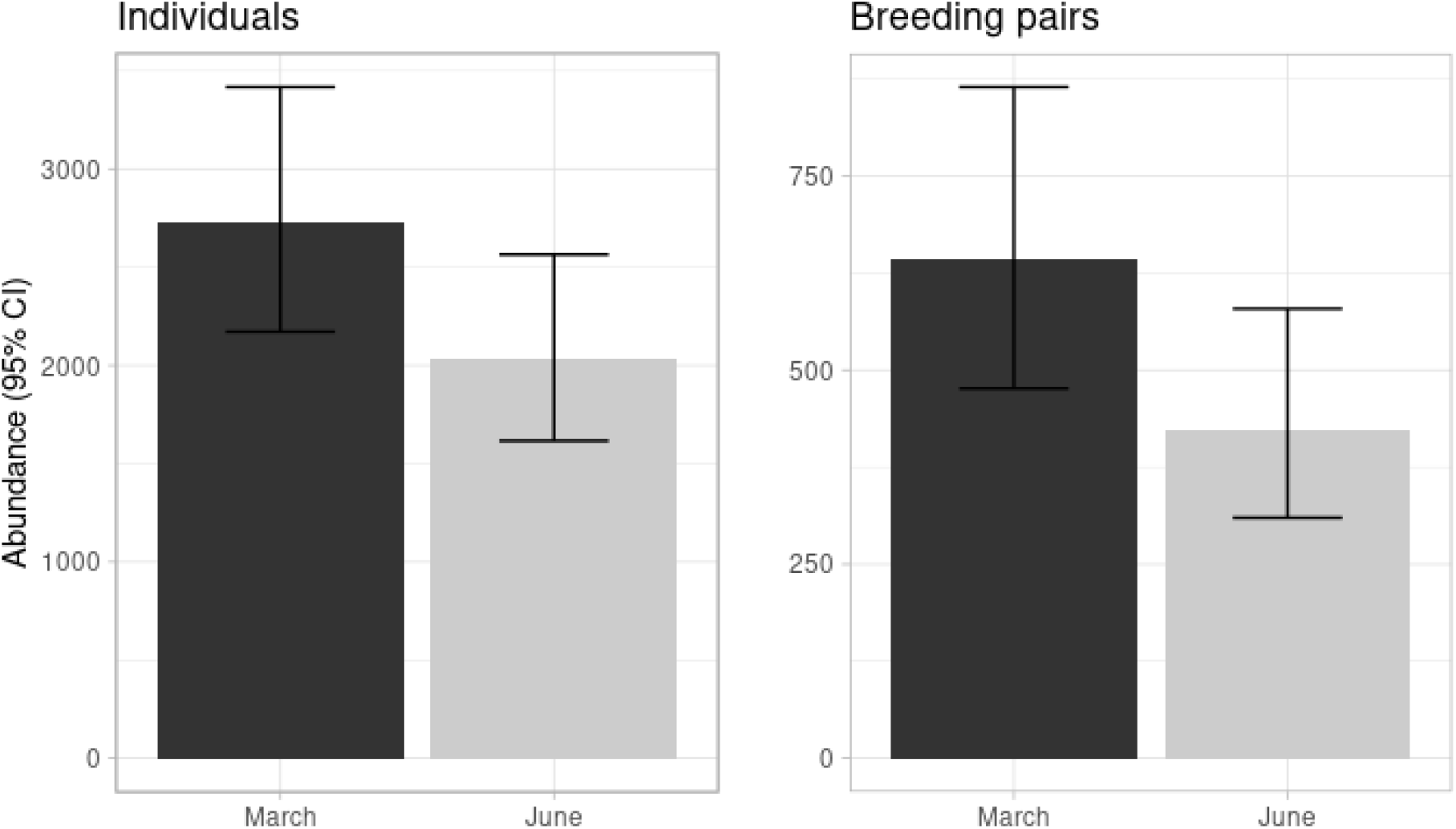
Estimated abundance ± 95% CI of individuals (left) and breeding pairs (right) of yellow-legged gulls in the historic centre of Venice in the considered months of 2018. Estimates were obtained by considering the models with the lowest AIC value (see Table 1)

The density and estimated number of breeding birds turned out to be less than a quarter of the estimated number of individuals (Table 2, Figure 2). The relative decrease of abundance recorded between spring and summer was slightly more pronounced for breeding pairs than for individuals (on average, −25% for individuals and −32% for breeding pairs), however, given the overlap of 95% CI, these estimates were not significantly different.

### Analysis of the initial effects of the new waste collection system on gull population

#### Comparison of the density estimation before (2017) and after (2018) the change of the waste collection policy

The fit of the models selected for the detection function of surveys performed in March and June of the years 2017 and 2018 from the selected eight Observation Points (OPs) was good both for individuals and breeding pairs after right-truncation (Table 3, Figures S3 and S4). However, as expected given the low number of OPs used in the analysis, the precision of density estimates obtained by the best models selected according to AIC was not very high (Table 4). In both years, the density of individuals and breeding pairs tended to be higher in March than in June (Table 4, Figure 3) with the estimate of individuals being quite comparable in the two years, while the one of breeding pairs markedly lower in 2018, both for March and June (Figure 3). These differences however were not statistically significant as the standard errors of the estimates were quite large and the 95% CI mostly overlapped, especially in March. Nevertheless, the decrease observed in June is noticeable, with the density recorded in 2018 corresponding about to the 66% of that recorded in 2017.

**Table 3.** Ranking of candidate models used for estimating yellow-legged gulls density in Dorsoduro and Santa Croce before (March and June 2017) and after (March and June 2018) the implementation of the new policy of waste collection. Best models were selected according to Akaike’s information criterion (AIC). K = Number of parameters; GOF (p) = p-value of the Cramér-von-Mises goodness of fit test; 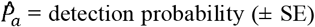; ΔAIC = difference in AIC from the best model.

| Survey | Model | K | GOF (p) | $\hat{P}_a \pm \text{SE}$ | ΔAIC |
| --- | --- | --- | --- | --- | --- |
| *Individuals* |  |  |  |  |  |
| March 2017 | Half-normal + Hermite polynomials | 1 | 0.84 | $0.29 \pm 0.02$ | 0.00 |
| Excluded observations: 23.2% | Half-normal + cosine | 1 | 0.84 | $0.29 \pm 0.02$ | 0.00 |
| | Hazard-rate + simple polynomial | 2 | 0.99 | $0.31 \pm 0.04$ | 2.35 |
| | Uniform + cosine | 2 | 0.96 | $0.28 \pm 0.03$ | 2.46 |
| June 2017 | Hazard-rate + simple polynomial | 2 | 0.86 | $0.26 \pm 0.05$ | 0.00 |
| Excluded observations: 18.5% | Uniform + cosine | 3 | 0.90 | $0.22 \pm 0.03$ | 2.60 |
| | Half-normal + cosine | 2 | 0.84 | $0.22 \pm 0.03$ | 3.12 |
| | Half-normal + Hermite polynomials | 1 | 0.05 | $0.35 \pm 0.03$ | 11.97 |
| March 2018 | Half-normal + Hermite polynomials | 1 | 0.94 | $0.33 \pm 0.02$ | 0.00 |
| Excluded observations: 19.2% | Half-normal + cosine | 1 | 0.94 | $0.33 \pm 0.02$ | 0.00 |
| | Uniform + cosine | 1 | 0.70 | $0.36 \pm 0.02$ | 0.73 |
| | Hazard-rate + simple polynomial | 2 | 0.90 | $0.36 \pm 0.04$ | 3.23 |
| June 2018 | Hazard-rate + simple polynomial | 2 | 0.98 | $0.31 \pm 0.06$ | 0.00 |
| Excluded observations: 19.3% | Uniform + cosine | 2 | 0.87 | $0.31 \pm 0.03$ | 1.90 |
| | Half-normal + cosine | 2 | 0.91 | $0.27 \pm 0.04$ | 2.94 |
| | Half-normal + Hermite polynomials | 1 | 0.16 | $0.39 \pm 0.03$ | 7.25 |
| *Breeding pairs* |  |  |  |  |  |
| March 2017 | Half-normal + cosine | 2 | 0.99 | $0.20 \pm 0.04$ | 0.00 |
| Excluded observations: 7.3% | Half-normal + Hermite polynomials | 1 | 0.41 | $0.29 \pm 0.03$ | 1.45 |
| | Uniform + cosine | 3 | 0.99 | $0.19 \pm 0.03$ | 1.74 |
| | Hazard-rate + simple polynomial | 2 | 0.97 | $0.25 \pm 0.05$ | 1.79 |
| June 2017 | Half-normal + cosine | 2 | 0.98 | $0.24 \pm 0.05$ | 0.00 |
| Excluded observations: 17.1% | Hazard-rate + simple polynomial | 2 | 0.91 | $0.24 \pm 0.11$ | 0.82 |
| | Uniform + cosine | 3 | 1.00 | $0.23 \pm 0.04$ | 0.95 |
| | Half-normal + Hermite polynomials | 1 | 0.23 | $0.42 \pm 0.05$ | 5.58 |
| March 2018 | Half-normal + Hermite polynomials | 1 | 0.89 | $0.29 \pm 0.03$ | 0.00 |
| Excluded observations: 12.7% | Half-normal + cosine | 1 | 0.89 | $0.29 \pm 0.03$ | 0.00 |
| | Uniform + cosine | 1 | 0.47 | $0.34 \pm 0.02$ | 0.82 |
| | Hazard-rate + simple polynomial | 2 | 0.74 | $0.35 \pm 0.05$ | 3.34 |
| June 2018 | Uniform + cosine | 1 | 0.81 | $0.42 \pm 0.04$ | 0.00 |
| Excluded observations: 13.3% | Half-normal + Hermite polynomials | 1 | 0.72 | $0.43 \pm 0.05$ | 1.85 |
| | Half-normal + cosine | 1 | 0.72 | $0.43 \pm 0.05$ | 1.85 |
| | Hazard-rate + simple polynomial | 3 | 0.86 | $0.38 \pm 0.11$ | 4.06 |

**Table 4.** Density of individuals and breeding pairs obtained for Dorsoduro and Santa Croce districts in both the surveyed years (2017 and 2018). Density estimates were calculated using the best models chosen according to the Akaike’s Information Criterion (AIC)

| **Survey** | **No. of observations**<br>**(clusters or pairs)** | **Density ± SE**<br>**(individuals or pairs/km<sup>2</sup>)** | **CV** |
| --- | --- | --- | --- |
| *Individuals* |  |  |  |
| March |  |  |  |
| 2017 | 188 | 348.9 ± 75.2 | 0.22 |
| 2018 | 251 | 352.4 ± 92.5 | 0.26 |
| June |  |  |  |
| 2017 | 207 | 274.1 ± 83.5 | 0.30 |
| 2018 | 251 | 284.8 ± 89.5 | 0.31 |
| *Breeding pairs* |  |  |  |
| March |  |  |  |
| 2017 | 101 | 104.9 ± 25.6 | 0.24 |
| 2018 | 124 | 87.6 ± 27.5 | 0.31 |
| June |  |  |  |
| 2017 | 107 | 92.4 ± 25.4 | 0.27 |
| 2018 | 123 | 60.9 ± 14.1 | 0.23 |

**Fig. 3.**
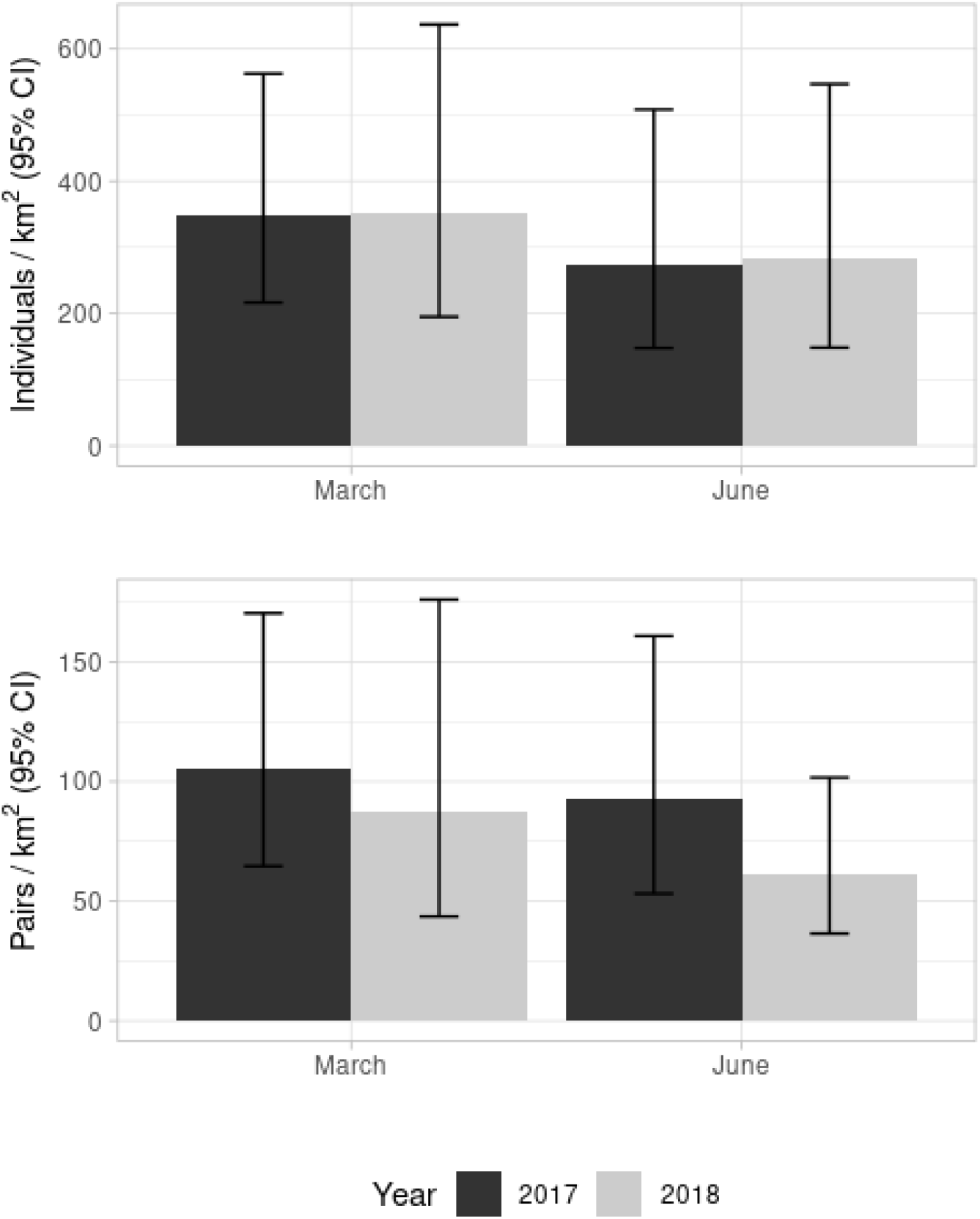
Estimated density ± 95% CI of individuals (above) and breeding pairs (below) of yellow-legged gulls in Dorsoduro and Santa Croce before (March and June 2017) and after (March and June 2018) the implementation of the new policy of waste collection. Estimates were obtained by considering the models with the lowest AIC value (see Table 3)

#### Analyses of the trend of yellow-legged gulls collected in Venice historical city centre by the service of wildlife recovery from 2010 to 2018

The GAM model fitting the trend of the number of yellow-legged gulls collected in the urban area by the service of wildlife recovery from 2010 to 2018 explained a large proportion of the deviance of the data (deviance explained = 92.2%, adjusted R^2^ = 0.90). As depicted in Figure 4, the two age classes considered for the analyses show a different trend, especially in the last three years. The number of >1-year birds increased over the sampling period at an increasing rate, while the number of 1-year birds together with *pulli* reached the peak in 2016 and then decreased markedly. In both cases, the fitted smooths turned out to be highly significant (>1-year birds: edf = 2.02, χ^2^ = 70.7, p ≪ 0.001; 1-year birds + *pulli*: edf = 3.52, χ^2^ = 36.2, p ≪ 0.001).

**Fig. 4.**
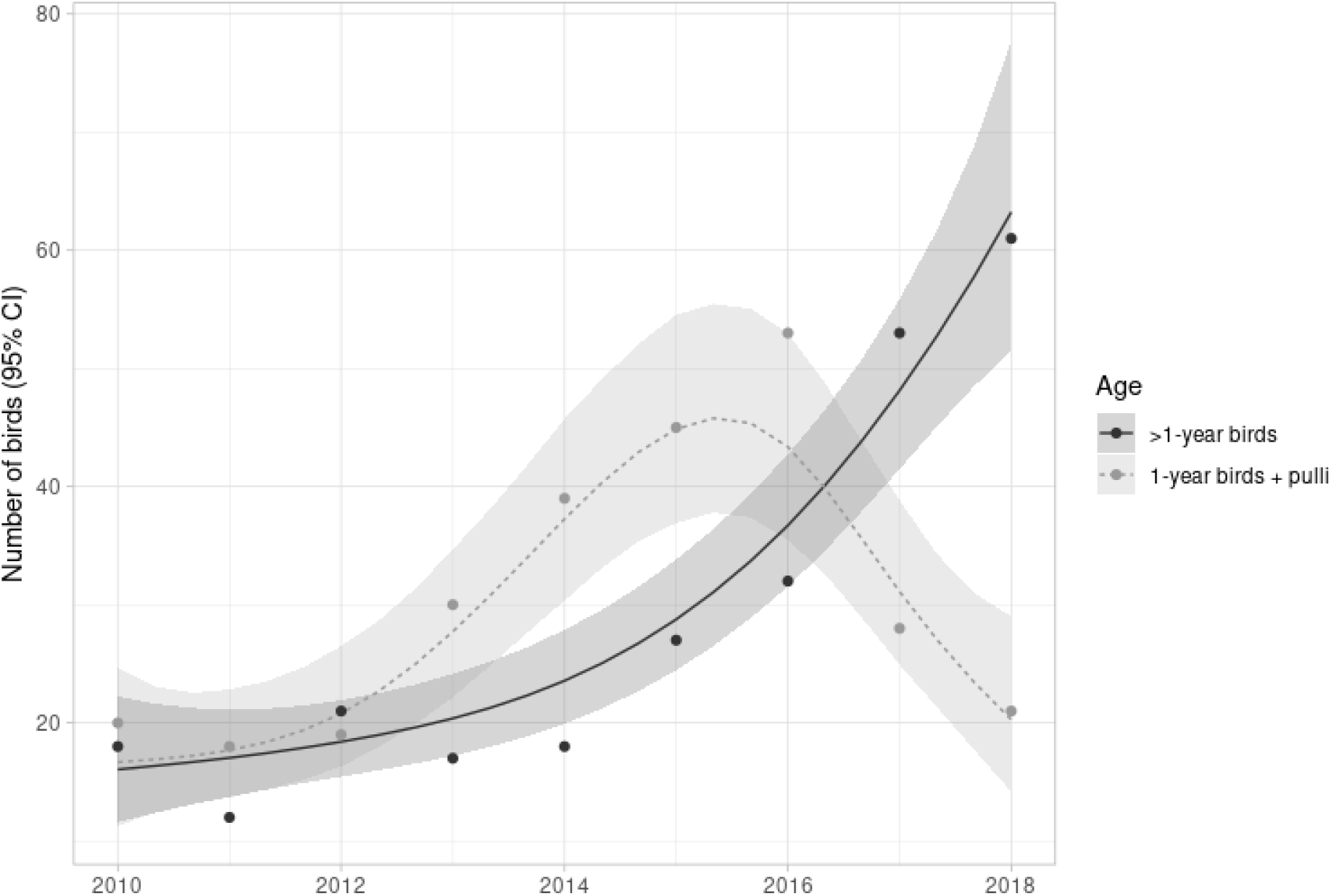
Number of yellow-legged gulls collected in Venice historical city centre by the service of wildlife recovery in the period 2010-2018. The fitted lines of the Generalized Additive Model with Poisson error distribution and logarithm as link function is also depicted for the two considered age classes (shaded area = 95% CI)

## Discussion

In this work we provided an updated estimate of the yellow-legged gull population in the historic centre of Venice and investigated the effects of the new waste collection system on the species by comparing the abundance of individuals and breeding pairs before and after the policy change, as well as by analysing the trend of yellow-legged gulls collected in the city centre by the wildlife recovery service over an eight-year period.

### Estimate of the abundance of yellow-legged gulls’ breeding population

For the estimation of the urban gull population we used the Distance Sampling method (Buckland et al. 2001). According to what is reported in literature (see Ross et al. 2016 for a review), this method has never been used before for census yellow-legged gulls in an urban environment. Most studies performed in urban settings used direct counts from vantage points and from street surveys (i.e. surveying rooftops from ground level) (Coulson and Coulson 2015) or conducted independently ground-based and aerial surveys (Durham 2003; Rock 2002; Sellers and Shackleton 2011). However, it has been proved that direct counts miss an appreciable number of nests, with an average maximum nest detection rates of 78% for vantage point surveys and 48% for street ones (Coulson and Coulson 2015), while aerial surveys lead to some issues in distinguishing between gull species (Durham 2003; Sellers and Shackleton 2011). Recently, the use of unmanned aerial vehicles (UAVs, or drones) for census of the urban-nesting gull population have also been used in the city of Victoria, Canada, with drones proving to readily discern the occupied nests and incubating birds that were undisturbed by them (Blight et al. 2019). However, flight restrictions for UAVs may render their use hard to practice in several urban settings (ENAC 2020).

Distance Sampling method has been successfully used in natural context to census breeding seabirds (Kirkwood et al. 2007; Lawton et al. 2006; Robertson et al. 2008). Moreover, its performance in estimating the total number of nests of a natural colony of large gulls has been compared with the strip transect counts method by Barbraud and colleagues (2014). Such study strongly advocates the use of distance sampling for surveying large gull colonies since estimates obtained by the strip transect count method were significantly lower (from 9 to 31%) than those obtained by distance sampling. These authors promote the use of distance sampling as it creates less disturbance than transect counts and it requires fewer observers and less time than direct counting methods (Barbraud et al. 2014). Therefore, it is reasonable to assume that the use of distance sampling is recommended also in urban environments, the latter being more complex than natural settings and with a higher risk of bias (Giunchi et al. 2007).

The density estimates of yellow-legged gull urban population in 2018 turned out to be quite reliable, given the precision and the consistency of the estimated detection probability among different models. Results indicated the presence of about 2700 individuals in March 2018 and 2000 in June, while the estimated breeding pairs were respectively about 600 and 400. The number of urban breeders therefore corresponds to approximately 45% (average value on the two estimates of March and June 2018) of estimated individuals. It is possible that some of the recorded individuals may use the city for foraging purposes only, while breeding elsewhere in the lagoon (Scarton and Valle 2017). Furthermore, it is known that an important fraction (between 15% and 30%) of the population of large gulls species can comprise non-breeding adults (Coulson et al. 1982; Kadlec and Drury 1968). In this case, the fraction is higher but the number of estimated individuals was maybe biased low as flying birds were not included in the counts.

Obtained estimates are rather different from those of 2005, when the urban colony was at the beginning of its growth and probably involved relatively young breeders (Soldatini et al. 2008), showing an average annual increase rate of about 25% between 2005 and 2018.

Trends similar to those shown by yellow-legged gulls in the historical centre of Venice have been found also in other urban areas, both in Italy (Benussi and Fraissinet 2020) and abroad (Rock 2013; Ross et al. 2016). As an example, in Britain and Ireland, large gulls breeding on rooftops showed an annual average increase rate of 17% for herring gulls and 28% for lesser black-backed gulls, *Larus fuscus*, between 1969 to 1976 (Monaghan and Coulson 1977). Both the species continued to increase until 1994, both in terms of number of urban breeding pairs and number of towns colonized (Raven and Coulson 1997), to reach in 1998-2002 the 20000 pairs of roof-nesting herring gulls, more than double the number recorded during the previous survey of 1994, and the 11000 pairs of lesser black-backed gulls, which is four times the number of 1994 (Mitchell et al. 2004). Estimates, however, are often obtained with different methods, posing a problem of comparability as well as a different degree of reliability of data.

Several factors have led to the colonization of urban areas by large gulls. First of all, the strong recovery experienced by many seabirds during the 20^th^ century after the decline of the 19^th^ century due to their persecution (Coulson 1963), and the consequent need of finding new sites suitable for breeding. Regarding this, buildings in towns are sort of man-made islands within a ‘sea’ of concrete that show many of the attributes afforded by natural breeding sites, as reported by Coulson and Coulson (2009). In the case of the Venice historical city centre, there are some peculiarities that make it particularly attractive to gulls, such as the massive presence of historical buildings and churches (n = 133 in the studied area, according to data provided by the Territorial Information Systems Service Office of the Venice municipality), all characterized by tiled roofs that make them inaccessible to people and therefore freely usable by yellow-legged gulls for nesting and resting purposes. Another characteristic of the city is the presence of a large number of street food, bars and restaurants providing take-away food that have exploded in the city in recent years to satisfy the ever-increasing demand for ‘hit and run’ tourism. The old town of Venice records over 10 million tourist presences every year to which are added the ‘visitor’ tourists (i.e. people which do not spend the night in the city), whose number is uncertain but, in any case, greater than a further 12 million/year (Campostrini and Dabalà 2017). The latter generally consume quick and cheap meals while strolling around the city, in this way attracting yellow-legged gulls that are waiting to steal their food or to eat the leftovers abandoned on the street (Coccon F. pers. obs.). Last, but not least, the city of Venice is surrounded by its lagoon that is the widest of the Mediterranean area, covering an area of 55000 ha, listed as Important Bird Area (Heath et al. 2000) and recognised as a Special Protection Area (SPA IT 3250046 Lagoon of Venice), according to the 147/2009 Birds Directive of the European Union. This synergy allows yellow-legged gulls to exploit either the resources of anthropogenic origin found in the city and the widespread natural sources offered by the lagoon for both nesting and feeding purposes. Importantly, the trend shown by the species in the Venice historical city centre is in contrast with that found in natural and artificial habitats (e.g. littoral strips, natural and artificial salt marshes, dredge islands, fish farms, restored habitats) of the north-eastern Adriatic coastline, where a moderate but statistically significant decline of the species has been recorded in the period 2008-2014 (Scarton et al. 2018). Additionally, if considering yellow-legged gulls breeding in the Venice lagoon, where most of nests are located in dredge islands and artificial salt marshes (Scarton and Valle 2017), an average annual increase rate of 1.3% was found between 2010 to 2018 (Scarton F. unpublished data), thus a fairly low value with respect to the one recorded in the old town. These findings are in line with those shown in previous work conducted on the west coast of Canada (Vermeer 1992), which found a higher annual increase rate of breeding glaucous-winged gulls, *Larus glaucescens*, in downtown Vancouver (9% yearly increase of roofs utilised by gulls and of number of gulls nesting on roofs), compared with that of the Strait of Georgia, where the average annual increase rate over the period 1975-86 was 2.6%. It is likely that the higher annual growth rate recorded in urban environment is linked to the higher reproductive success due to the lower predation rate than in natural habitats (Coulson and Coulson 2009; Monaghan 1979), besides the larger availability of food, in the form of domestic and commercial waste and leftovers, which are easily reachable also by inexperienced juveniles (Luniak 2004).

Our results show a decreasing tendency of the yellow-legged gulls abundance from March to June. This finding is in line with that found in Rome, where the sharp decline recorded by the urban population of the species in summer months has been interpreted as an abandonment of the city for searching for food in other sites, following the significant decrease of people staying in the city during holidays and therefore of trophic resources available to gulls (Fraticelli 2017). However, this explanation is not plausible for the city of Venice, where the tourism rate is high all over the year and actually increases in the summer period (Città di Venezia 2018). In this case, the decrease of the yellow-legged gull urban population observed in June is probably linked to the abandonment of the city by those individuals that have failed their breeding attempt(s) and thus move elsewhere in search of attractive sources, present both in the lagoon and on the mainland (Coccon et al. 2015), to reduce intraspecific competition within the city.

### Analysis of the initial effects of the new waste collection system on gull population

Our study did not show any decrease of individuals between 2017 and 2018 but highlighted a decrease of breeding pairs which was particularly evident in June (−34%). Such a decrease is not statistically significant, probably because of the small number of observation points (n = 8) used for the comparison of the population between the two years that leads to a low precision of the estimates. This limitation, however, is unsurpassable since for 2017 we had data available for two districts only, due to unavailability of funds. Anyway, this result is particularly encouraging as it predicts a decline of new-born gulls in the city. It also strengthens the thesis supported by Belant (1997) that limiting availability of food, in the case of Venice by preventing the rubbish accumulation in the street, is a key element in controlling the urban population of the species since it reduces the overall suitability of the area for gulls. These findings agree with previous work (Pons 1992) according to which a reduction in trophic availability tends to have a much more dramatic effect on breeding success of the species than it does on adult survival, as adults are able to adapt and find other forms of sustenance (Pierotti and Annett 1991). Interestingly, results provided by distance sampling are in line with those obtained by analysis of data from the wildlife recovery service. Indeed, the number of *pulli* and 1-year birds recovered showed a clear decline after 2016, i.e. the year of implementation of the new waste collection system in a portion of the city. It is possible that this trend is actually due to a decrease of breeding pairs, as suggested by Distance Sampling, which led to a reduction in the number of chicks born, and to limited food availability following the policy change (Coccon and Fano 2020), which possibly forced juveniles to move elsewhere in search of new foraging areas, as competition for trophic resources probably increased in the city. On the contrary, the number of recovered >1-year birds showed a noticeable increase after 2016. It might be supposed that this increase may be due to the enhanced competition for limited resources that may have led animals to fight for grabbing food, therefore to injure themselves (F. Coccon pers. obs.), or to ingest worse quality food or even starving, with consequent higher probability of finding them in difficulty.

To sum up, the available data suggest that the new waste management policy affect not only the number of foraging yellow-legged gulls, as reported by Coccon & Fano (2020), but also the abundance of breeding pairs and possibly their breeding success, even though our data does not provide a clear-cut demonstration of this effect. This result is likely affected by the short period (i.e. two years) considered in this analysis and a clearer pattern will probably emerge in the coming years.

## Conclusions

In this study, we updated the estimate of the urban population of yellow-legged gulls for the historic centre of Venice using, Distance Sampling (Buckland et al. 2001), a novel monitoring method for the species in urban environment. This method provided a reasonable estimate of population size, outlining a picture very far from the last one described for the area by Soldatini and Mainardi (2006). This underlines the importance of monitoring, to be carried out possibly continuously or at least every 2-3 years, to detect numerical and behavioural changes in the species and provide an updated picture of the urban population in order to properly manage it. There are, however, some intrinsic limits in the estimates obtained in this work. In fact, it is important to note that the estimate of individual abundance is probably biased low as we did not consider birds in flight. Future studies should be focused on performing a new estimate of the urban population of yellow-legged gulls to evaluate the long-term effects of the new waste collection policy. New estimates may be improved by considering both flying and roosting animals (Bibby et al. 2000). Furthermore, it would be interesting to study the effects on the target species of measures to combat and contain the Covid-19 pandemic which has profoundly changed the city, by emptying it from tourists and daily visitors. This would indirectly provide information on the influence of tourism on the presence and distribution of the species in the city.

## Acknowledgments

We thank L. Panzarin for assistance in data collection and the Venice Municipalized Company Veritas Spa for financially supporting the project. We are particularly grateful to AVM S.p.A. – Azienda Veneziana della Mobilità, T Fondaco Dei Tedeschi by DFS, Venezia Terminal Passeggeri S.p.A. and the priestly Church of Venice for the opportunity of access to bell towers and the other elevated observation points used for monitoring. A special thank goes to R. C. Jones for reviewing and improving the English language.

