## Supplementary material for "More than expected: the abundance of yellow-legged gulls *Larus michahellis* breeding in the historic centre of Venice and the initial effects of the new waste collection policy on the population"

Manuscript submitted to ‘Urban ecosystems’ Journal

**Table S1** Number of residents, official B&B, urban waste production and surface area for each of the six districts of the historical centre of Venice. Data regarding number of residents and B&B were provided by the Statistics and Research Office of the Municipality of Venice (data updated to 1^st^ January 2017), while data on daily waste production were provided by the public waste management company of Venice, Veritas (data updated to 31th December 2017). Districts’ extension were calculated using the GIS platform (ESRI, ArcGIS 10.2 for Desktop).

| District | Inhabitants (No. residents) | B&B (No. beds) | Urban waste production (Tons/day) | Area (km^2^) |
| --- | --- | --- | --- | --- |
| Cannaregio | 15605 | 301 | 40 | 1.40 |
| Castello | 13424 | 318 | 32 | 1.90 |
| Dorsoduro | 6430 | 95 | 28 | 0.97 |
| Santa Croce | 4939 | 169 | 16 | 1.42 |
| San Polo | 4612 | 100 | 16 | 0.34 |
| San Marco | 3750 | 199 | 32 | 0.54 |

**Table S2** Observation points (OP) used for monitoring yellow-legged gulls. The building type, height and survey effort (i.e. the proportion of urbanized area included in the buffer calculated at 500 m) are reported here. With regards to the bell towers, the height refers to the top, while in the case of the Patriarchal Seminary (OP 3), the building of the Maritime Port (OP 4) and the private one (OP 8) it refers to the terrace floor. The heights of the bell towers were obtained from the works of Urbani De Gheltof (1892) and Sammartini and Resini (2002) the remaining ones were provided by the responsible persons and owners of the buildings.

| OP | OP Name | OP Buiding type | OP Height (m) | Survey Effort |
| --- | --- | --- | --- | --- |
| 1 | Church of Carmini | Bell tower | 66 | 0.87 |
| 2 | Church of S. Trovaso | Bell tower | 53 | 0.70 |
| 3 | Patriarchal Seminary of the Basilica of Madonna della Salute | Panoramic terrace | 27 | 0.50 |
| 4 | Unused building of the Tronchetto Maritime Port | Roof top | 15 | 0.48 |
| 5 | Municipal Garage of the Piazzale Roma bus terminal | Roof top | 25 | 1.00 |
| 6 | Church of S. Nicolò dei Mendicoli | Bell tower | 26 | 0.70 |
| 7 | Church of S. Cassiano | Bell tower | 43 | 1.00 |
| 8 | Private building | Panoramic terrace | 20 | 1.00 |
| 9 | Church of S. Geremia | Bell tower | 43 | 1.00 |
| 10 | Church of Madonna dell’Orto | Bell tower | 56 | 0.57 |
| 11 | Fondaco dei Tedeschi Building | Terrace | 20 | 1.00 |
| 12 | Church of S. Francesco della Vigna | Bell tower | 69 | 0.65 |
| 13 | Church of S. Giuseppe di Castello | Bell tower | 22 | 0.80 |
| 14 | Church of S. Elena | Bell tower | 52 | 0.44 |
| 15 | Basilica of S. Maria Gloriosa dei Frari | Bell tower | 69 | 1.00 |
| 16 | Basilica of S. Marco | Bell tower | 97 | 0.72 |

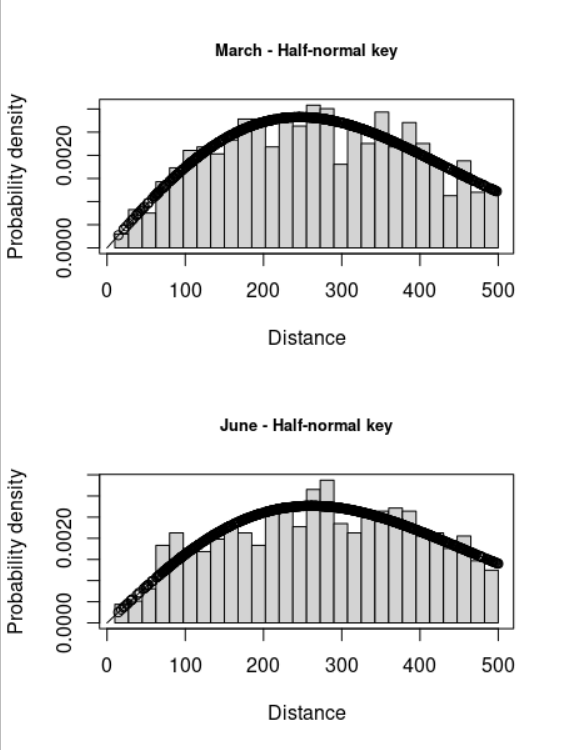

**Fig. S1** Histograms of distances of yellow-legged gulls (i.e. number of individuals) recorded in March and June 2018 in the historic centre of Venice from the selected 16 vantage points (OPs). The probability density functions of the detection probability estimated using the best model selected by the Akaike Information Criterion, AIC, is overlaid in each plot

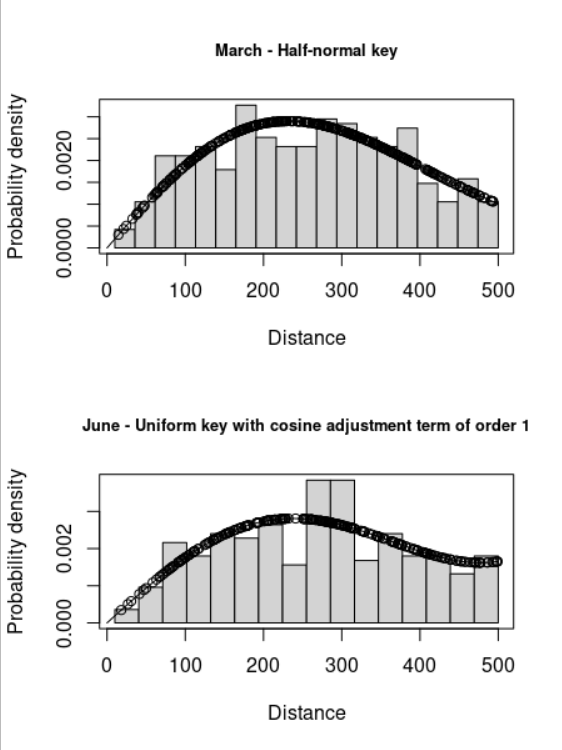

**Fig. S2** Histograms of distances of yellow-legged gulls (i.e. breeding pairs) recorded in March and June 2018 in the historic centre of Venice from the selected 16 vantage points (OPs). The probability density functions of the detection probability estimated using the best model selected by the Akaike Information Criterion, AIC, is overlaid in each plot

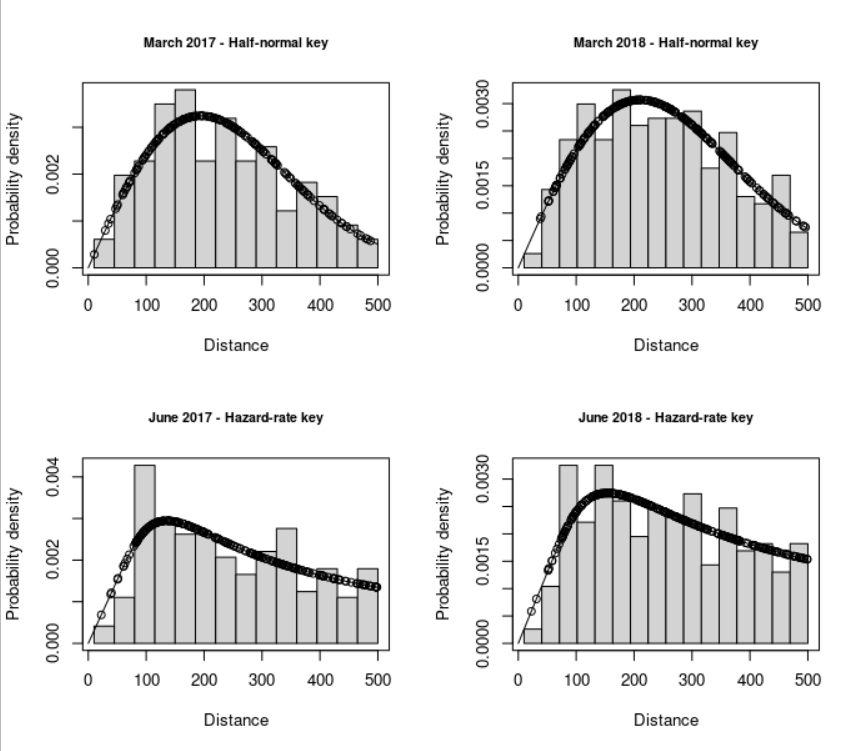

**Fig. S3** Histograms of distances of yellow-legged gulls (i.e. number of individuals) recorded in Dorsoduro and Santa Croce before (March and June 2017) and after (March and June 2018) the implementation of the new policy of waste collection from eight Observation Points (OPs). The probability density functions of the detection probability estimated using the best model selected by the Akaike Information Criterion, AIC, is overlaid in each plot

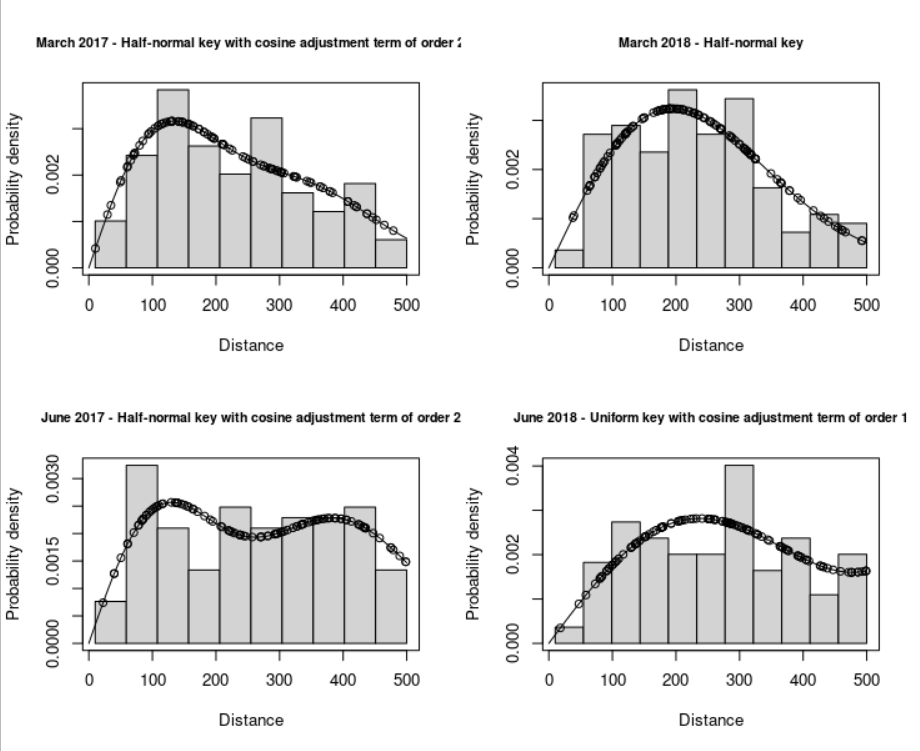

**Fig. S4** Histograms of distances of yellow-legged gulls (i.e. breeding pairs) recorded in Dorsoduro and Santa Croce before (March and June 2017) and after (March and June 2018) the implementation of the new policy of waste collection from eight Observation Points (OPs). The probability density functions of the detection probability estimated using the best model selected by the Akaike Information Criterion, AIC, is overlaid in each plot

Urbani De Gheltof GM (1892) Venezia dall'alto. Filippi Editore, Venezia
